# Genome-wide copy number variation-, validation- and screening study implicates a novel copy number polymorphism associated with suicide attempts in major depressive disorder

**DOI:** 10.1101/534909

**Authors:** Shitao Rao, Mai Shi, Xinyu Han, Marco Ho Bun Lam, Guangming Liu, Yun Kwok Wing, Hon-Cheong So, Mary Miu Yee Waye

**Affiliations:** The Nethersole School of Nursing, The Croucher Laboratory for Human Genomics, The Chinese University of Hong Kong, N.T., Hong Kong SAR, China; Department of Psychiatry, The Chinese University of Hong Kong, N.T., Hong Kong SAR, China; School of Biomedical Sciences, Faculty of Medicine, The Chinese University of Hong Kong, N.T., Hong Kong SAR, China; College of Food and Biological Engineering, Jimei University, Xiamen, Fujian, China; KIZ-CUHK Joint Laboratory of Bioresources and Molecular Research of Common Diseases, Kunming Institute of Zoology and The Chinese University of Hong Kong, China; CUHK Shenzhen Research Institute, Shenzhen, China

## Abstract

**Background:** The genetic basis of suicide attempts (SA) remained unclear, especially for the copy number variations (CNVs) involved. The present study aimed to identify the susceptibility variants associated with SA among major depressive disorder (MDD) patients in Chinese, covering both single-nucleotide polymorphisms and CNVs.

**Methods:** We conducted GWAS on MDD patients with or without SA and top results were tested in a replication study. A genome-wide CNV study was performed. Subsequently, a validation assay using the qRT-PCR technology was performed to confirm the existence of the associated CNV and then applied to the entire cohort to examine the association.

**Results:** In CNV analysis, we found that the global rate of CNV was higher in SA compared to non-SA subjects (p=0.023). The genome-wide CNV study revealed a SA-associated CNV region that achieved genome-wide significance (corrected p-value=0.014). The associated CNV was successfully validated and identified to be a common variant in this cohort and its deletion rate was higher in suicide attempters (OR=2.05). Based on the GTEx database, genetic variants that probe this CNV was significantly associated with the expression level of *ZNF33B* in two brain regions (p-value<4.2e-05). Besides, there was a significant interaction between neuroticism and the CNV in affecting suicidal risk; the CNV showed a significant effect (OR=2.58) in subjects with high neuroticism only.

**Conclusions:** We identified a new common CNV that may be involved in the etiology of SA. These findings imply an important role of common CNVs in the etiology of SA, which suggests a new promising avenue for investigating the genetic architecture of SA.

## 1. Introduction

An estimated 10 to 20 million of people attempted suicide yearly resulting in both acute injuries and long-term disability all over the world (Bertolote and Fleischmann, 2002; World_Health_Organization, 2012). History of suicide attempts (SA) is also the most accurate predictor of suicide completion, which is the 10^th^ leading cause of mortality worldwide (Hawton and van Heeringen, 2009; Chang et al., 2011; Varnik, 2012). Moreover, suicide represents the 2^nd^ leading cause of death in the 14-24 age groups in Europe (OECD, 2012). The number of suicide victims in China has been estimated to account for around 42% of suicide death in the world (Phillips et al., 2002; Weiyuan, 2009). Suicide is believed to be a public health issue that is affected by a combination of numerous biological, psychological, social and environmental factors (Hawton and van Heeringen, 2009; Sher, 2011; Beghi et al., 2013). Genetic studies, such as family, twin and adoption, consistently suggested the involvement of genetic factors in the etiology of SA (Roy et al., 1991; Brent et al., 1996; Pedersen and Fiske, 2010). With around 90% of SA subjects having at least one kind of psychiatric disorders, the predisposition to SA is partly dependent on the presence of psychiatric disorders, especially major depressive disorder (MDD) (Brent et al., 1993; Henriksson et al., 1993; Suominen et al., 1996). However, the occurrence of SA is believed to be partly independent of these disorders as it is also affected by psychological correlates, e.g. impulsivity, aggression and conscientiousness domain (Brent et al., 1996; Brent and Mann, 2005; Brezo et al., 2006; Rao et al., 2018).

With substantial advances in high-throughput technologies, it is now possible to investigate the genetic architecture of SA at a whole-genome level, e.g. by genome-wide SNP arrays and whole genome sequencing. Compared with the conventional candidate-gene association studies, the whole-genome approaches could screen hundreds of thousands of variants simultaneously (Shi et al., 2011). From the classic SNP-by-SNP analysis in GWAS, most studies however did not find strong evidence at genome-wide significant association or only with a marginal association (reviewed by Sokolowski *et al.*) (Sokolowski et al., 2014). Although two previous genome-wide association studies of SA in major depression reported several genome-wide significant or suggestive signals, both of the studies were performed in European populations, and the heritability explained by the reported loci were small (Perlis et al., 2010; Schosser et al., 2011). An alternative approach to investigating the genetic etiology of SA is to perform genome-wide copy number variation (CNV) study. Although CNVs has no greater intrinsic pathogenic potential than SNPs, their size could let them potentially influence the related gene product at each intersecting gene or change the genomic context with *cis*- or *trans*-effects (Perlis et al., 2012). CNVs have already been found to be associated with many psychiatric disorders, including autism spectrum disorder, schizophrenia, bipolar disorder and MDD (Szatmari et al., 2007; International Schizophrenia, 2008; Ingason et al., 2011; Malhotra et al., 2011; O’Dushlaine et al., 2014).

Genome-wide CNV studies in suicidal behavior has begun several years ago but no conclusive results has been reported, not even for one validated CNV (Perlis et al., 2012; Gross et al., 2015; Sokolowski et al., 2016b). Perlis and colleagues identified SA-associated CNVs from 189 major depressive patients with SA and 1,073 major depressive patients without SA in the sequenced treatment alternatives to relieve depression (STAR*D) clinical trial (Perlis et al., 2012). They mainly focused on the CNVs with a size >100 kb and a frequency <10% but failed to find any CNV that could reach genome-wide significance. Gross *et al.* performed a genome-wide CNV study for 475 cases with suicidal behavior (199 SAs and 276 suicides) and 1,133 controls without suicidal behavior (Gross et al., 2015). The initial algorithms Penn-CNV identified two potential CNVs that were only present in cases but not in any control, however neither the visual inspection of the raw data nor the following validation assay using the qRT-PCR technology could support this finding. Besides, Sokolowski and colleagues adopted family-based trio samples (n= 660 trios) with severe SA in offsprings to conduct a CNV-based GWAS using a calling of rare (<1%) and large (>100 kb) CNVs (Sokolowski et al., 2016b). The study did not find any significantly associated CNV. Although some rare CNVs were found in a subset of SA, they elaborated that these CNVs were not further confirmed with any validation method. The cases recruited in Gross *et al.* and Sokolowski *et al.* were not limited to patients with a history of MDD.

In summary, the genetic basis of SA, especially for the associated CNVs, remained poorly understood. Here we conducted the first genome-wide CNV study of SA in Chinese in order to identify the potential SA-associated CNVs. For the genome-wide significant CNV regions and intersecting genes we not only validated them by the quantitative RT-PCR (qRT-PCR) technology, but also determine their distributions in suicide attempters in a larger cohort of MDD subjects. Furthermore, interaction between the associated CNVs and the assessed psychological correlates on risk of SA was also investigated. On the other hand, we also carried out the first GWAS of SA in major depression in a Chinese population, trying to uncover further susceptibility SNPs. The signals around the genome-wide suggestive threshold were replicated in an independent Chinese cohort and then confirmed by meta-analyses. Notably, a recent study reported that the genetic correlation (r_g_) of MDD in Europeans and Chinese is only moderate (r_g_ of lifetime MDD= 0.33), suggesting some genetic risk factors underlying depression may not be common across populations (Bigdeli et al., 2017). The same may also apply for the genetic basis of suicidality in MDD.

## 2. Materials and Methods

### 2.1 Participants

In this study, we recruited 122 MDD patients with suicide attempts (SA, 27 men and 95 women) and 175 MDD patients without suicide attempts (non-SA, 56 men and 119 women). The inclusion and exclusion criteria have been described in our previous studies (Rao et al., 2016; Rao et al., 2018). For each subject, a lifetime SA was identified in a semi-structured diagnostic interview by an experienced clinical psychiatrist. The psychological, social and occupational functioning of each subject was rated by the Global Assessment of Functioning (GAF) scale, which is described on page 34 in the text revision of the Diagnostic and Statistical Manual of Mental Disorders, 4th edition (DSM-IV-TR). Besides, the Chinese-Bilingual Structured Clinical Interview for the Diagnostic and Statistical Manual of Mental Disorders (Axis I, Patient version) (CB-SCID-I/P) was also carried out to ascertain if the subject has any kind of psychiatric disorders. In addition, all the recruited subjects were restricted to be Han Chinese in origin and above 18 years old. Those subjects who suffered from mental retardation or dementia were excluded from this study. Written informed consents were obtained from all the subjects after completely explaining the purpose and process of this study. This study was reviewed and approved by the Joint Chinese University of Hong Kong - New Territories East Cluster Clinical Research Ethics Committee (Reference Number: *CRE-2006.393*).

### 2.2 Assessment of subject’s psychological correlates

Psychological correlates of subjects were assessed by NEO Personality Five-factor Inventory (NEO-FFI), Barratt Impulsiveness Scale (BIS) and Hospital Anxiety and Depression Scale (HADS). The Chinese version of the three scales had been translated, validated and demonstrated to have a good agreement with the English version (Leung et al., 1999; Yang et al., 1999; Yao et al., 2007). Briefly, the NEO-FFI is a simplified version of NEO Personality Inventory Revised (NEO PI-R). It scores individuals on five personality dimensions: NEO-O, NEO-C, NEO-E, NEO-A and NEO-N (Eggert et al., 2007). In Hong Kong, Cheung and colleagues had constructed a normative data based on a cohort of 1,421 Hong Kong Chinese healthy college students (Cheung et al., 2008). Subjects in this study were characterized by converting their domain *T-scores* into *z-scores* based on the Chinese normative data. Subsequently, we could classify the *z-scores* of the five personality dimensions into low, average or high levels respectively. Participants whose *z-scores* were at least one-half standard deviation (SD) below or above the average were classified into a low or high group respectively. The others were classified into an average group. Besides, BIS is a self-report measurement of impulsiveness. A recent review suggested that the intent of subject’s impulsiveness could be classified into 3 levels, i.e. high, normal limits and low level of impulsiveness (Stanford et al., 2009). Moreover, an optimal balance between sensitivity and specificity of HADS could be achieved if the case was defined by a score of 8 or above for both the anxiety and depression subscales (Bjelland et al., 2002).

### 2.3 Pilot genotyping assays in SNP-array, following replication studies and meta-analysis

Genomic DNA samples of all subjects were extracted from 2 ml of saliva reagent and then measured for concentration by Nanodrop 2000c spectrophotometer. A subgroup of subjects comprising 48 SA and 48 non-SA were chosen for the pilot genome-wide genotyping assay in Illumina iScan system using the HumanOmni ZhongHua-8 v1.3 DNA Analysis BeadChip Kit (Illumina, Inc.) at the Li Ka Shing Institute of Health Sciences of the Chinese University of Hong Kong. The raw data generated from the system were transformed in the Illumina GenomeStudio program (Illumina, Inc.) and then used for further statistical analyses.

Stringent quality controls were firstly adapted for the SNP-array data to exclude low-quality subjects and variants in PLINK v1.9 program (Purcell et al., 2007). Briefly, subjects with missing data rates ≥10% were dropped from the analysis, and single nucleotide polymorphisms (SNPs) were dropped from the analysis if they had either a minor allele frequency <1%, a missing data rate ≥1% or a Hardy-Weinberg equilibrium (HWE) p-value <10e-04. Finally, a cleaned sample set consists of 86 major depressive patients (50% SA) and 745,680 SNPs remained for further association studies.

After quality control, genome-wide association study (GWAS) between the qualified SNPs and SA was conducted. The likelihood ratio p-values were reported. The conventional p<5e-08 and p<5e-06 were considered as the genome-wide significant and suggestive thresholds respectively (Dudbridge and Gusnanto, 2008; Ioannidis et al., 2009). The top GWAS signals around the suggestive threshold were then selected for the following replication studies in an independent subgroup of major depressive patients consisted of 74 SA and 127 non-SA subjects. The TaqMan^®^ SNP genotyping assays were employed for the replication experiments in QuantStudio™ 7 Flex or ViiA 7 Real-Time PCR system (Life Technology, USA). The PCR mixture and program were the same as the descriptions in our previous study (Rao et al., 2016). A fixed-effects meta-analysis of the results from both the pilot GWAS and the following replication study was performed to determine the association between the top SNP signals and SA in the combined sample set (117 SA and 170 non-SA). Besides, gene-based tests were performed using the Functional Mapping and Annotation (FUMA) tool with the Eastern Asian reference panel population from the 1000 genome project Phase III database (Watanabe et al., 2017). Since approximately 15,000 protein coding genes were involved in this test, the gene-based association significant and suggestive threshold was set at p<3.33e-06 (0.05/15,000) and p<3.33e-05 (one order higher than the genome-wide threshold) respectively.

### 2.4 CNV detection from the SNP-array data, following validation assays and further screening assays

The raw data generated from the iScan Illumina system was analyzed in the GenomeStudio program to detect CNVs with a CNV analysis algorithm (cnvPartition 3.2.0, Illumina), which combined the Log R ratio and the B allele frequency to determine the copies (Seiser and Innocenti, 2014). It can also work with GenomeStudio’s GenomeViewer to visualize copy number regions graphically. Before the CNV detection, we firstly conducted quality control, in which subjects with missing data rates ≥10% were dropped from the analysis. A total of 10 subjects were removed, resulting in 86 subjects (50% SA) for CNV calling. Only CNVs with a confidence threshold ≥35 and a minimum probe count ≥3 were called out for further analyses. Burden analyses of segmental CNVs in terms of differences in rate, proportions, total length and average length between SA and non-SA MDD patients were carried out. Deleted and duplicated CNVs were also extracted separately for burden analyses. In addition, association analyses of each segmental CNV with SA were carried out in the overall group as well as in the subgroups of the deleted and duplicated CNVs. To identify if there is any gene associated with SA, gene-based association tests were performed for a set of 18,870 known genes.

We note that CNV calling based on SNP arrays is subject to error and should not be regarded as confirmatory (Marenne et al., 2011; Xu et al., 2013). We consider CNV calling from the GWAS SNP-array as a first step to screen for possible CNV regions, yet the results will be experimentally confirmed with more reliable assays.

With regard to the validation of significantly associated CNVs or genes, subjects harboring such CNVs or genes and a sample of randomly chosen controls were selected for the following validation assays. For the associated segmental CNV, 3 customized Taqman probes were designed to validate the existence of the CNV. For the detailed information of the three probes please refer to Supplementary Figure 1. Briefly, the physical positions of the three SNP probes within the 10q11.21 region (the CNV region detected by the calling algorithm) in the SNP-array were identified. Based on their location, we custom-made three corresponding CNV probes aiming at validating the existence of the associated CNVs. In view of positive results in the 1st and 2nd probes, we further screened the whole sample (122 SA and 175 non-SA) so that we could identify the frequency of the CNV in SA and non-SA subjects. To test for the existence of CNV(s) intersecting with the *ZNF33B* gene (a gene prioritized from statistical analysis, see later sections), we ordered one pre-designed probe (Hs04385285_cn) from Thermo Fisher Scientific. The CNV probe located in the *RNase P* gene was employed as a reference assay because this gene is stable in human species and invariably exists as 2 copies.

The qRT-PCR assay was run on the QuantStudio™ 7 Flex Real-Time PCR System (Life Technology, USA). The PCR mixture was prepared and the PCR program was set according to the manufacturer’s instructions. Briefly, 10 μl PCR mixture contains 2 μl of genomic DNA per subject, 5 μl of TaqPath™ ProAmp™ Master Mix, 0.5μl of each 20× Taqman probe (target and reference probes) and 2 μl of deionized H_2_O. The mixture was firstly treated by activation of the uracil-N-glycosylase for 2 min at 50 °C, followed by denaturation for 10 min at 95 °C and amplification over 40 cycles of 15 s at 95 °C and 1 min at 60 °C. Once finished, all data was exported from the system and then analyzed in the CopyCaller software (v2.0). Each subject was run in triplicate. Moreover, every plate takes two stable subjects as reference samples in order to keep the consistence of calling CNV between different plates. If the obtained results of subjects were different between the SNP-array data and the validation assay, the subjects would be probed again.

Comparisons of continuous and categorical variables were accomplished by t test and Chi square test respectively using SPSS (version 20.0, IBM Analytics). Fisher’s exact test was adopted when a low cell of observed or expected frequencies occurred in the analyses. The interaction effect between the associated CNV and the neuroticism domain was assessed by the logistic regression analysis after adjustment for sex and age.

## 3. Results

### 3.1 Demographics and psychological correlates of SA and non-SA

A total of 122 SA and 175 non-SA were recruited in this study. The two groups are sex- and age-matched subjects (Table 1, p-value = 0.062 and 0.570 respectively). A low employment rate was observed in SA (37.7%) and the rate was significantly lower than that in non-SA (p-value = 0.025). Additionally, the global functioning of the suicide attempters was significantly lower than that of the non-suicide attempters (Table 1, GAF, p-value = 0.001). With regard to the psychological correlates of subjects, suicide attempters were found to have higher score on neuroticism than the non-suicide attempters (Figure 1, p-value = 0.023). However, the other correlates did not demonstrate any significant difference between the two groups (Figure 1, p-value > 0.081).

**Figure 1.**
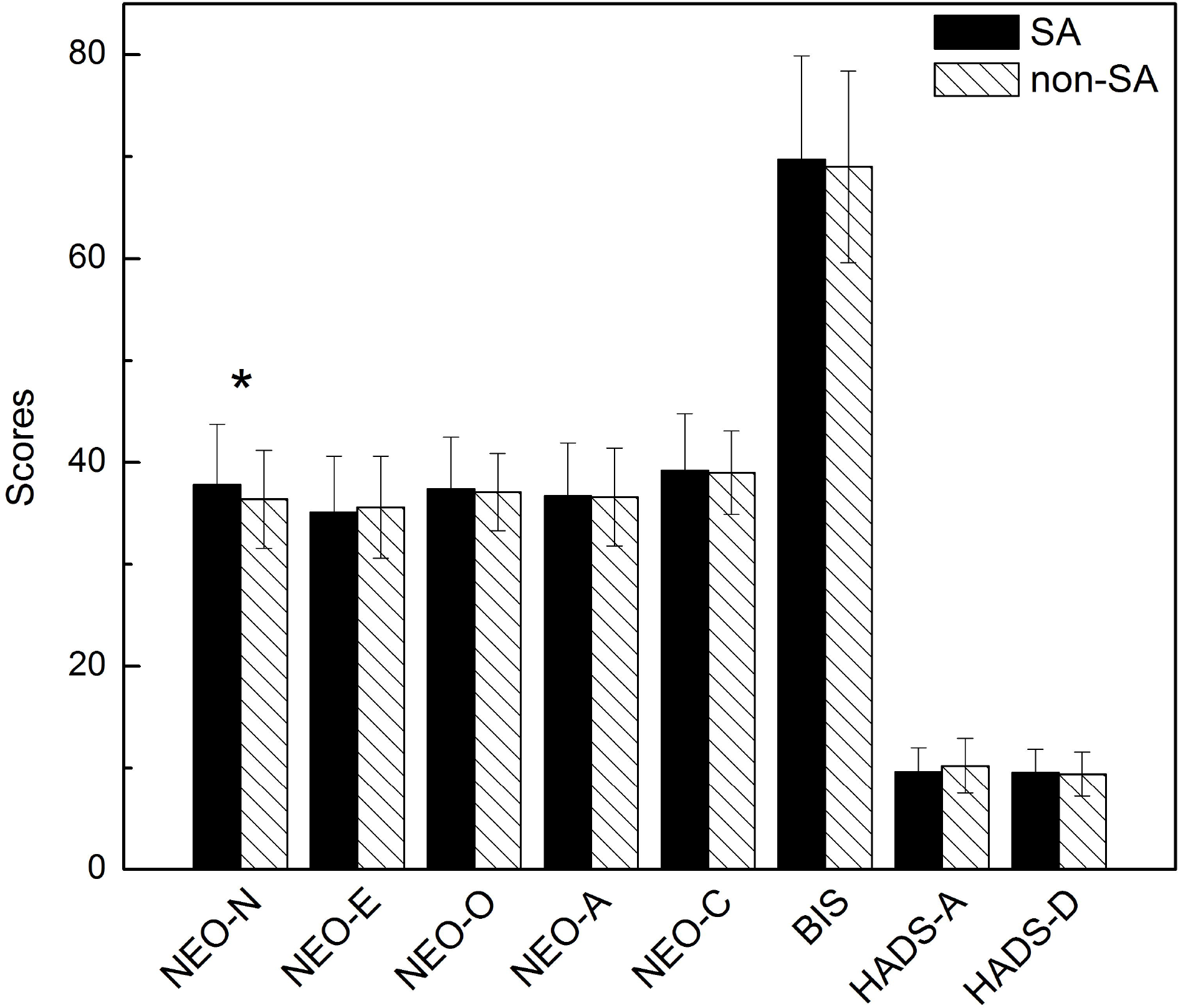
Comparison of psychological correlates between SA and non-SA. NEO-N, NEO-E, NEO-O, NEO-A and NEO-C: the five personality dimensions assessed by the NEO-FFI; BIS: the severity of impulsiveness assessed by the Barratt Impulsiveness Scale; HADS-A or HADS-D: Depression or Anxiety subscale in the Hospital Anxiety and Depression Scale. *P-value <0.05.

**Table 1.** Demographics of MDD patients with SA (SA, n=122) or without SA.

| Phenotype | SA<br>(n=122) | non-SA<br>(n=175) | Statistics | P-value |
| --- | --- | --- | --- | --- |
| Female (%) | 95(77.9) | 119(68.0) | $X^2 = 3.48$ | 0.062 |
| Age, mean ±SD | 42.6 ± 11.9 | 43.4 ± 12.2 | t = -0.57 | 0.570 |
| Employment (%) | 46(37.7) | 89(50.8) | $X^2 = 5.02$ | **0.025** |
| GAF, mean ±SD | 45.7 ± 19.7 | 53.3 ± 16.4 | t = -3.44 | **0.001** |

MDD: subjects with major depressive disorder; SA: MDD patients with suicide attempts; non-SA: MDD patients without suicide attempts; GAF: score of Global Assessment of Functioning. P-value below 0.05 is in bold.

### 3.2 Pilot genome-wide association studies of SA, following replication studies and meta-analyses

A subgroup of 48 SA and 48 non-SA was chosen for the pilot GWAS. After stringent quality controls, 43 SA and 43 non-SA remained for further association analyses. The cleaned SA and non-SA are also sex- and age-matched subjects (Table S1, p-value for differences in sex and age distribution= 0.486 and 0.710 respectively). From the pilot GWAS, a total of 8 top SNP signals were around the suggestive threshold for association (p<5e-06) (Manhattan plot shown in Figure S2A). Besides, there are 53 signals at the p<1e-05 level from GWAS (Table S2). Table 1 highlights the 8 top signals and there are 2 SNPs (rs4863321 and rs3844582) having an initial p-value below the suggestive threshold (p-value = 2.83e-06 and 3.54e-06 respectively). Moreover, four of the eight signals are located closely on a chromosome 11 region (Figure S2B, circled by blue) and found to be in strong LD (r^2^=1, please also refer to the LD matrix in Figure S3). One of the four SNPs (rs10895780), together with the other four SNPs in table 1, were chosen for replication studies carried out in an independent cohort comprising 74 SA and 127 non-SA subjects. As shown in Table 2, only one SNP (rs3844582) showed a marginal association with SA (Table 2, replication p-value = 5.79e-02), but the association could not withstand correction for multiple testing (p= 0.05/5= 0.01). Subsequently, we performed a meta-analysis using the results from both the pilot GWAS and the replication studies. The SNP rs3844582 had the lowest combined p-value (Table 2, p = 5.09e-05).

**Table 2.**
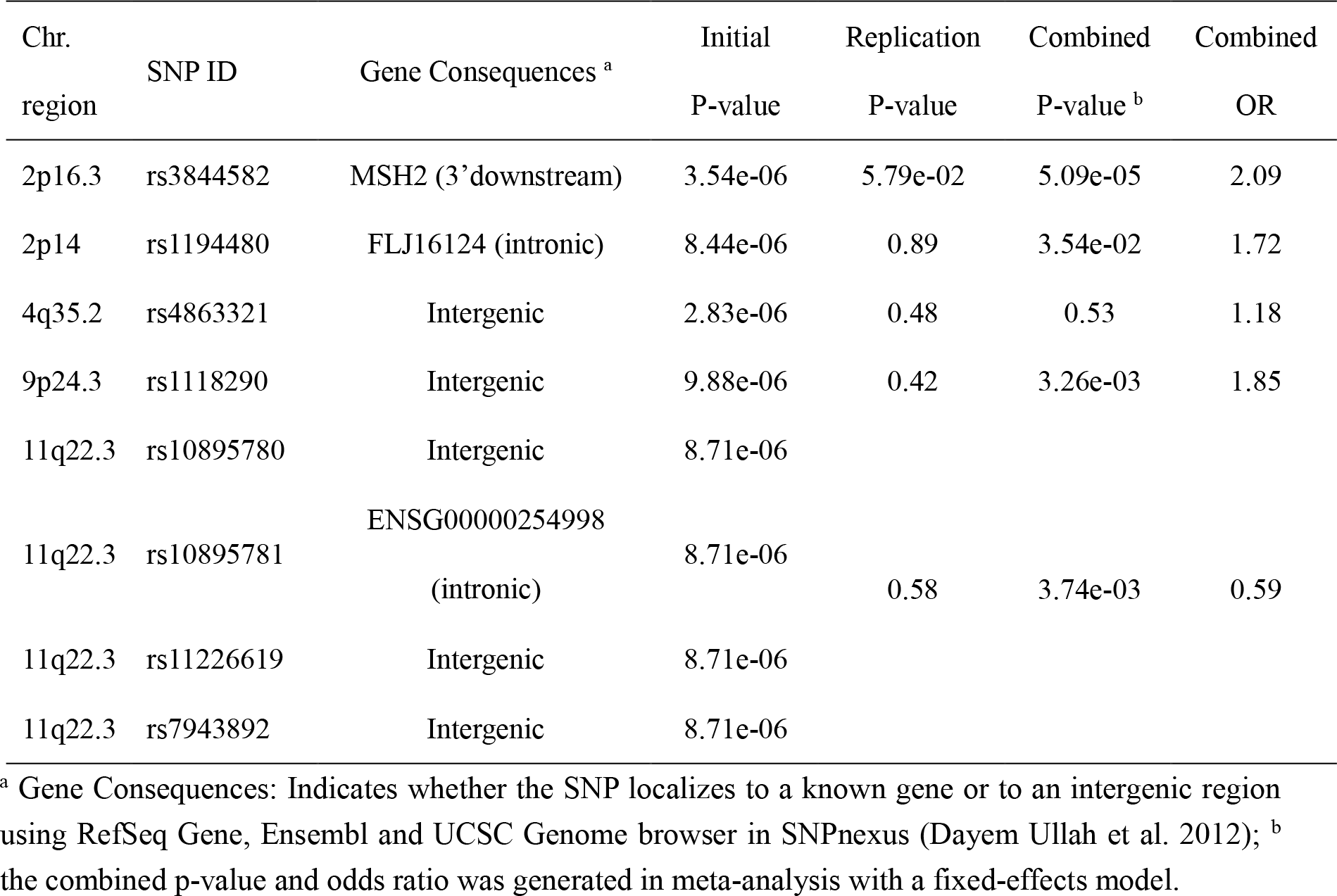
Top signals associated with SA from the pilot GWAS.

We then performed gene-based association tests in which all the input SNPs were mapped to around 15,000 protein coding genes. The most significant results were observed for the *TPH2* gene (z-score = 3.95, p-value = 3.93e-05) and the *WDR82* gene (z-score = 3.74, p-value = 9.27e-05) (Figure S4). The TPH2 protein, representing the isoform II of the rate-limiting enzyme in the biosynthesis of serotonin, was widely reported to be associated with various psychiatric disorders and suicidality in candidate case-control association studies (Zill et al., 2004; Lopez de Lara et al., 2007; Gonzalez-Castro et al., 2014). However, none of the 6 previously GWAS reported significant association of the *TPH2* gene with SA, even for a nominal association (Perlis et al., 2010; Schosser et al., 2011; Willour et al., 2012; Mullins et al., 2014; Galfalvy et al., 2015; Sokolowski et al., 2016a). Nevertheless, some of above GWAS did not restrict the subjects to MDD patients and all of them were based on Caucasian samples; heterogeneity in the samples may partially explain differences in study results.

### 3.3 Burden analysis and association analyses of SA with segmental CNVs and a set of known genes respectively

From the cleaned 43 SA and 43 non-SA subjects, we detected a total of 3,146 CNVs from the SNP-array data. Firstly, we conducted burden analysis for the overall group (Figure 2). We observed that the global rate of CNV (any type) was 1.12 times higher in SA subjects compared to the non-SA group (p-value = 0.023). In addition to analyzing the overall burden of CNV, deletion and duplication CNVs were also extracted separately for burden analysis. As shown in Figure 2, there were significant differences in global rate and total length in the duplicated CNVs (1.96 times compared to non-SA group, p-value = 0.008 and 1.90 times compared to non-SA group, p-value = 0.005 respectively). However, we did not observe any significant differences with regards to deletion CNVs (p-value > 0.496).

**Figure 2.**
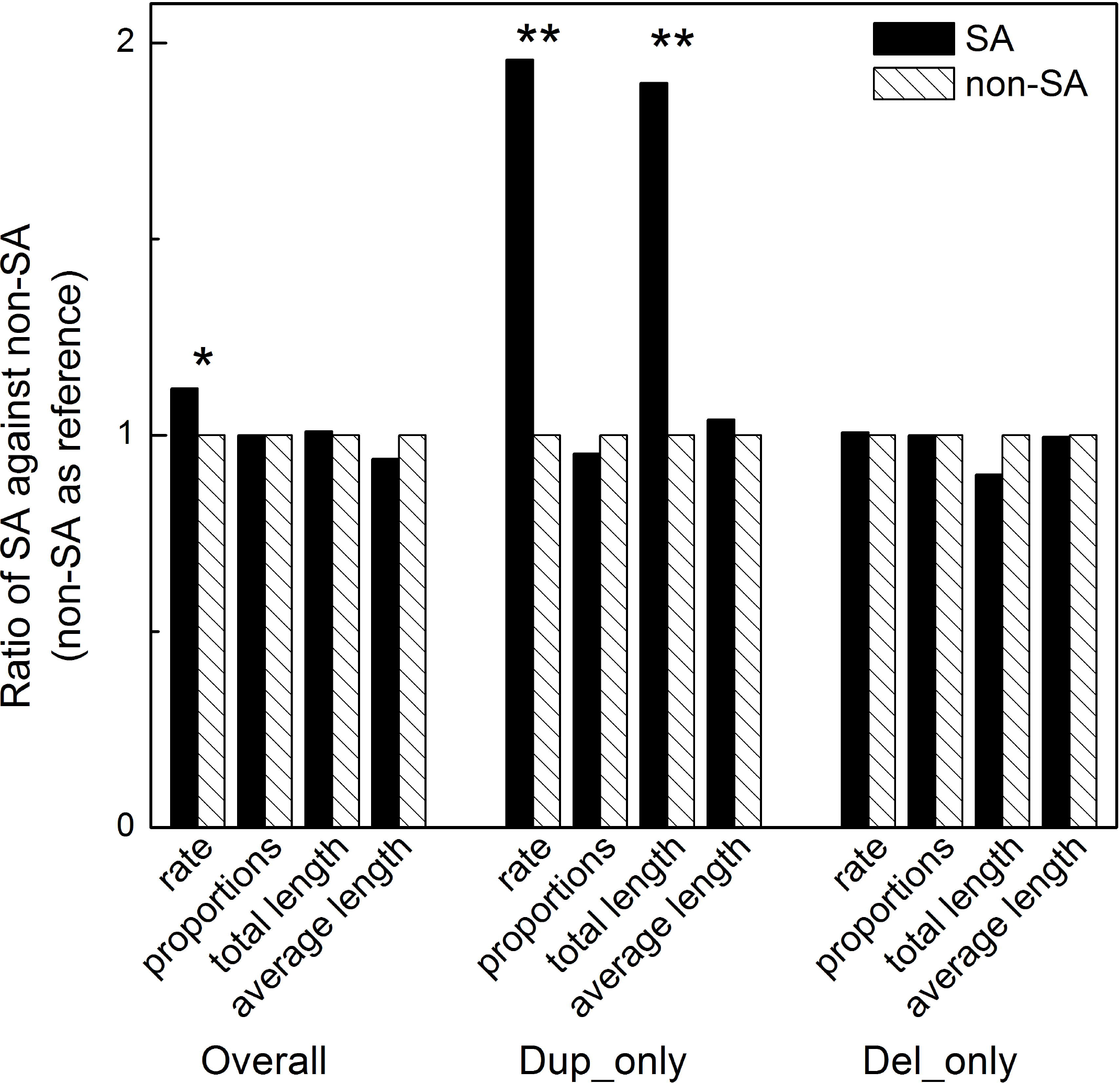
Burden analysis of segmental CNV for overall group, deletion and duplication subgroups respectively. Rate: rate of segmental CNVs in group; proportions: proportion of samples with one or more segments; total length: total length spanned; average length: average segment size. SA and non-SA: MDD patients with or without SA. * P-value < 0.05, ** p-value < 0.01.

We also carried out association analyses of each segmental CNV with SA in the three groups. In the overall group, association analysis did not yield any CNV segment that was significantly different between the SA and non-SA (Table 3, genome-wide corrected p-value > 0.176). Association analysis in the duplicated subgroup, however, identified a CNV segment that could withstand the genome-wide correction (Table 3, corrected p-value = 0.014) and present in 9 of the 43 suicide attempters but not in any of the 43 non-suicide attempters. Visual inspection of the raw data in GenomeViewer program supports the presence of the CNV (Figure S5, marked by blue arrows). With regard to the gene-based analysis (we examined the genes intersecting with the CNVs) with SA in the overall group, association analysis did not observe any known gene that could withstand the correction of 18,870 tests (Table 4, p-value > 0.102). Interestingly, all the top 5 associated genes belong to the zinc-finger protein family. In the duplication subgroup, the *ZNF33B* gene located in the 10q11.21 region could withstand the correction of multiple testing (Table 4, corrected p-value = 0.010), but the visual inspection of the raw data did not support this finding (Figure S5, marked by red arrows). As remarked earlier, CNV calling from the SNP-array was considered as a screening step mainly for prioritization of interesting regions in follow-up assays. Also, the copy number of CNVs quantified by the computer algorithm may not be accurate. A previous study showed a low sensitivity (0.2-0.3) but high specificity (0.97-0.99) for CNV-calling algorithms on the Illumina arrays (Marenne et al., 2011); the positive predictive value by the cnvPartition algorithm (which is also used in this study) is ~0.83 (Figure 4 in the reference), suggesting that it is worthwhile to follow up the top results in more reliable experimental assays.

**Table 3.** Associations of segmental CNVs with SA in the overall group, the duplication subgroup and the deletion subgroup respectively.

| *Group/CNV<sup>a</sup>* | *Position<sup>b</sup>* | *Number of*<br>*SA subjects* | *Number of*<br>*non-SA subjects* | *Nominal*<br>*p-value* | *Corrected*<br>*p-value<sup>c</sup>* |
| --- | --- | --- | --- | --- | --- |
| ***Overall*** |  |  |  |  |  |
| 10q11.21 Dup | 42406669..42661203 | 9 | 0 | **5.0×10<sup>-4</sup>** | 0.176 |
| 19q11 Dup | 27811892..28220337 | 12 | 3 | **0.011** | 0.763 |
| 6p22.1 Del | 28019979..28022248 | 6 | 0 | **0.012** | 0.899 |
| ***Dup_only<sup>d</sup>*** |  |  |  |  |  |
| 10q11.21 Dup | 42406669..42661203 | 9 | 0 | **1.2×10<sup>-3</sup>** | **0.014** |
| 19q11 Dup | 27811892..28220337 | 12 | 3 | **0.010** | 0.093 |
| 3p11.1 Dup | 90310715..90498372 | 6 | 0 | **0.013** | 0.139 |
| ***Del_only<sup>e</sup>*** |  |  |  |  |  |
| 19p13.3 | 417262..427262 | 3 | 0 | 0.118 | 0.955 |
| Xp22.33 | 3218815..3218815 | 7 | 3 | 0.158 | 1 |
| 9p24.3 | 2028942..2029454 | 2 | 0 | 0.244 | 1 |

<sup>a</sup> CNV: Only the top 3 signals were present here. <sup>b</sup> Positions are in base pairs for UCSC Build hg19. <sup>c</sup> corrected p-value: genome-wide corrected p-value. <sup>d</sup> Dup\_only: the analysis was performed within the duplication subgroup. <sup>e</sup> Del\_only: the analysis was performed within the deletion subgroup. Abbreviations: SA: suicide attempts; non-SA: non-suicide attempts. P-value below 0.05 are in bold.

**Table 4.**
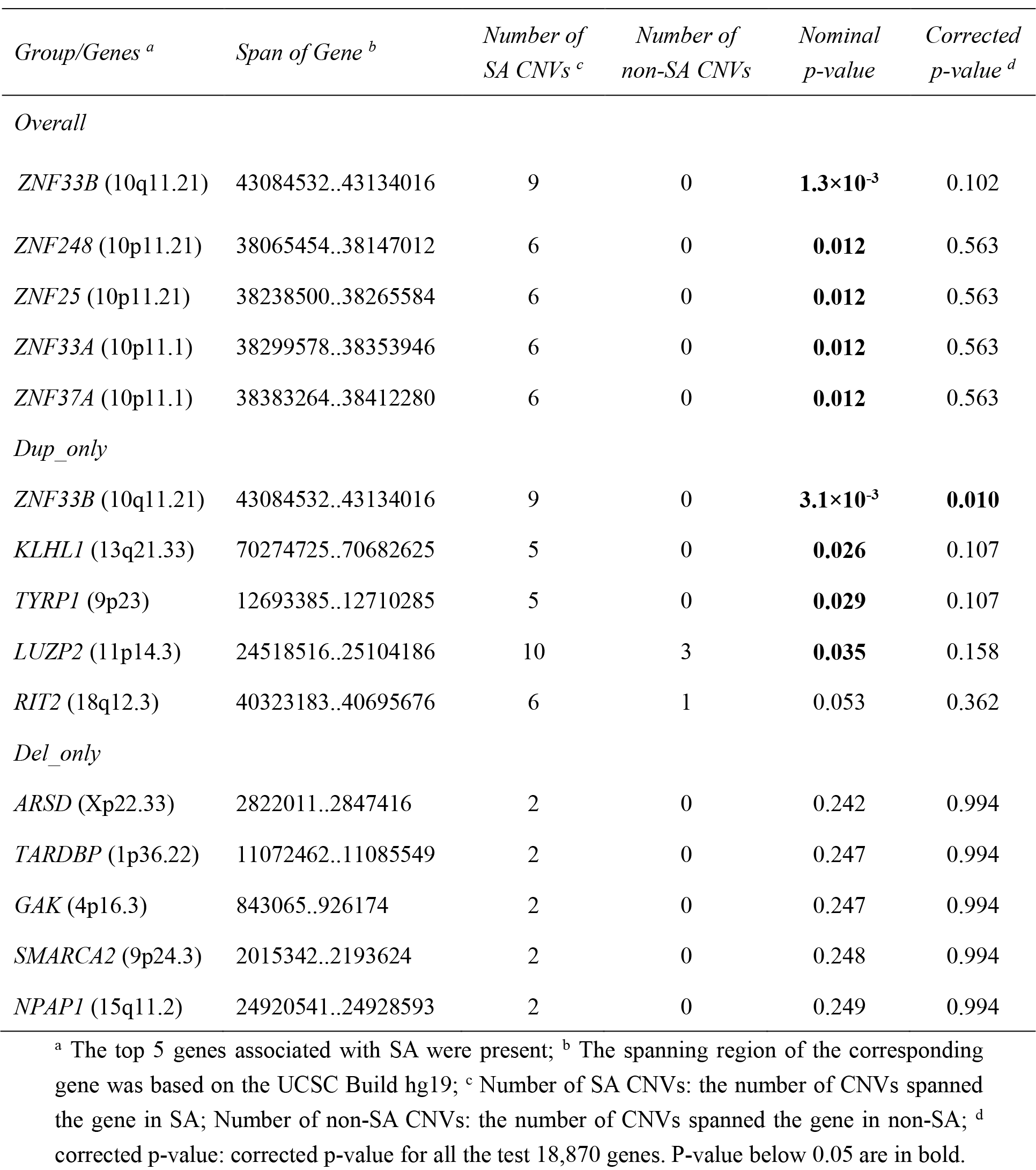
Gene-based associations of segmental CNVs with SA in the overall group, the duplication subgroup and the deletion subgroup respectively.

Before further experimental validation, we first searched for more detailed information of the probes within the 10q11.21 region in our SNP-array panel. We found that there were 3 probes spanning approximately 117kb (Figure S1, 41,716,457-41,833,540, hg38). The 3 probes correspond to 3 SNPs respectively (rs80241791, rs75876199 and rs79507803). The former two SNPs were found to be significantly associated with the expression level of the *ZNF33B* gene in a total of 14 tissues in the Genotype-Tissue Expression (GTEx) database, including two brain regions (pituitary, p-value = 2.7e-05, normalized effect size = 0.41; cerebellum, p-value = 4.2e-05, normalized effect size = 0.60).

### 3.4 Validation assays for the presence of the associated CNV and the *ZNF33B* gene

For validating the presence of the associated CNV observed from the SNP-array data, we custom-made three CNV detection kits based on the detailed information of the 3 probes within the 10q11.21 (Figure S1). The 9 SA subjects with the associated CNV and an equal number of non-SA subjects were chosen for the validation assays. As shown in Figure S6, two subjects have duplicated copies and one subject has a deleted copy in the 1st probe (Figure S6 left). As for the 2nd probe, the same two subjects with duplications in the 1st probe have duplicated copies (Figure S6 right). However, we did not observe any variations in the 18 SA or non-SA subjects for the 3rd probe (Figure S7). Besides, we also ordered one pre-designed probe for validating the presence of the CNV intersected with the *ZNF33B* gene but did not find variation in any of the subjects (Figure S8). Although we cannot detect CNVs intersecting with *ZNF33B*, it is still possible that CNVs outside the gene may regulate gene expression of *ZNF33B*.

Our findings suggest that CNV-calling algorithms for GWAS may be used to locate and prioritize the regions of potentially associated CNVs, but are far from perfect to determine the accurate CNV copy numbers. Follow-up validation assays are required to confirm the copy numbers. As the validated CNV may only have a small to moderate effect on suicidal risk, testing 18 subjects is clearly not adequate. We therefore decided to follow up this candidate CNV in all the recruited patients (N = 297) with the same experimental assay.

### 3.5 Customized Taqman copy number probes for CNV detection in the whole sample and evaluating interaction with neuroticism

Given the findings above, we conducted qRT-PCR assays with our customized Taqman copy number probes for the whole sample (122 SA and 175 non-SA), including the 96 subjects included in the original SNP-array. The validated CNV was found to be common in subjects and could be considered as a copy number polymorphism (CNP). As illustrated in Figure 3, the frequency of the CNP in the 2nd probe in SA subjects was much higher than that in non-SA subjects (adjusted p-value = 0.066, OR = 1.74) in the entire sample, although the difference only showed a trend towards significance. However, there was indeed a significant difference between cases and controls if we only calculated the deletion rate (Figure 3, adjusted p-value = 0.045, OR = 2.05), while the duplication rate did not have a significant difference (adjusted p-value = 0.663). With regard to the 1st probe, we did not observe any significant difference in the whole sample, deletion or duplication rate (Figure 3, p-value > 0.490).

**Figure 3.**
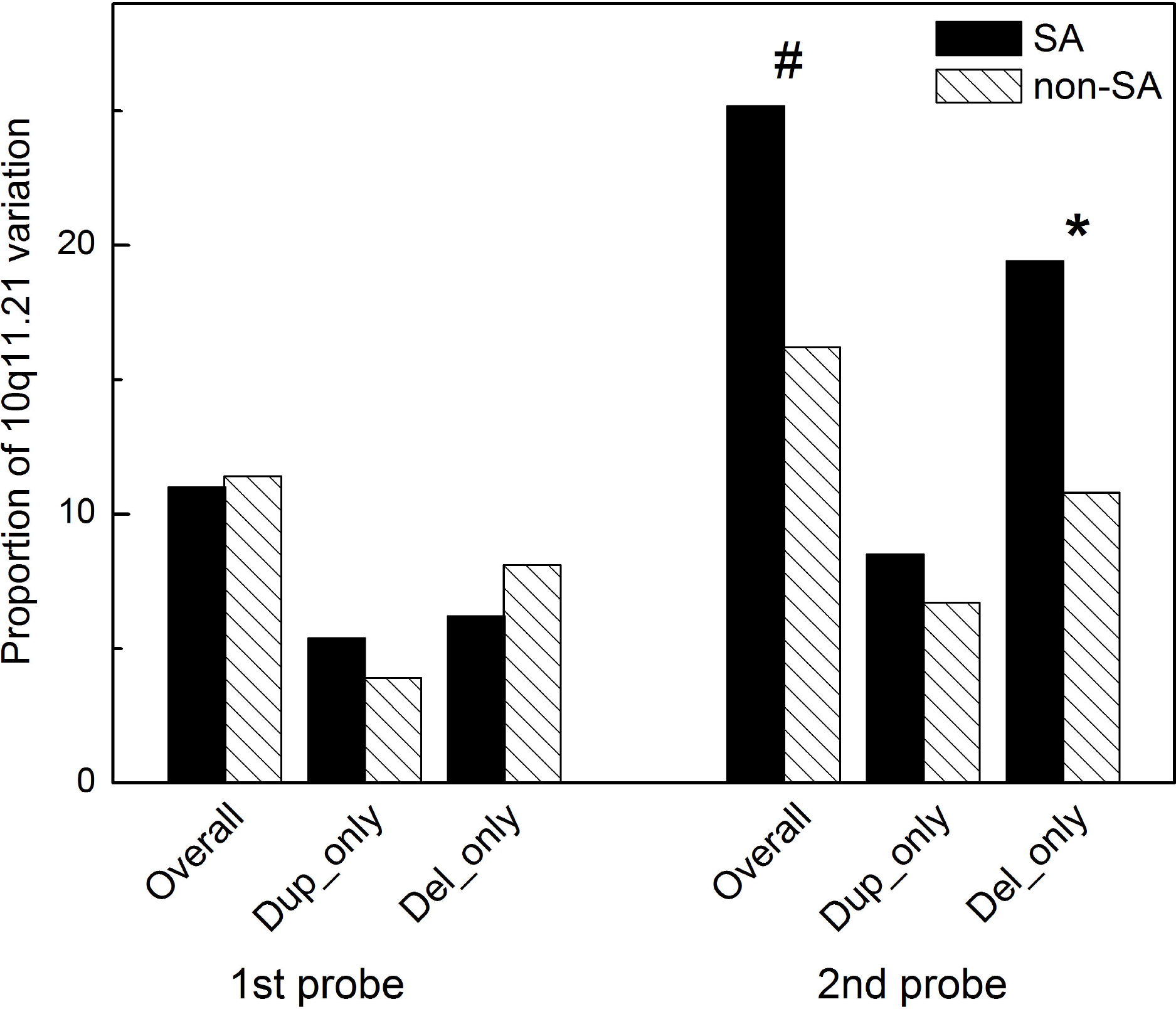
Frequency of the associated CNP between SA (n=122) and non-SA (n=175) with two customized Taqman copy number probes. P values and odds ratio was generated in logistic regression analysis after adjustment of sex and age. * P-value < 0.05; ^#^ P-value < 0.1.

**Figure 4.**
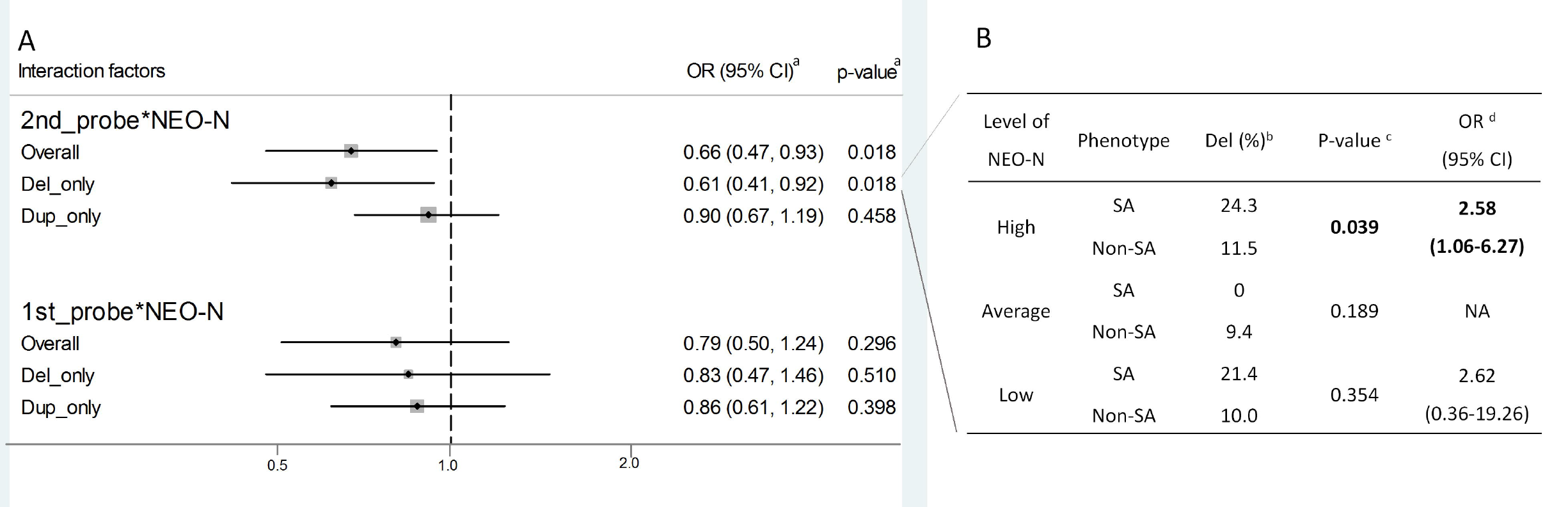
Evaluation of interaction effects between the neuroticism domain and the CNP in 1st probe and 2nd probe respectively on the risk of SA. ^a^ P values and odds ratio was generated in logistic regression analysis after adjustment of sex and age; ^b^ Only the deletion rate of the CNP was compared between SA and non-SA subjects; ^c^ P values were generated by the Chi square test; ^d^ Odds ratio was generated in logistic regression analysis controlling for sex and age in the stratified subgroup.

We then tested for interaction between neuroticism and the identified CNP with suicide. We tested 2 models in which neuroticism was included as a continuous trait or a categorical trait. We observed that the continuous neuroticism score significantly interacted with the candidate CNP as detected by the 2nd probe (p-value for interaction= 0.031 and 0.024 respectively when we considered overall association and deletion only).

We also transformed the continuous scores of the neuroticism domain into categories (high, average or low) according to the constructed normative data by Cheung and colleagues (Cheung et al., 2008). Notably, the CNP as detected by the 2nd probe showed significant interaction with neuroticism categories (Figure 4A, both adjusted p-values = 0.018).

To further explore the interaction effect, we conducted separate association studies of the CNP with SA in subjects with high, average or low levels of neuroticism (Figure 4B). A greater genetic effect was observed in the subgroup of subjects with a high level of neuroticism (Figure 4B, p-value = 0.039, OR = 2.58). On the other hand, the association either in the average or low level subgroup did not reach significance (Figure 4B, p-value > 0.189).

## 4. Discussion

In the present study, a new CNP within 10q11.21 was found to be associated with SA in major depression. The CNP region was first prioritized from a genome-wide CNV study and further validated by an experimental assay. We then performed the assay on all recruited depression subjects, and found that deletion of the CNP was associated with higher suicidal risks (adjusted p-value = 0.045, OR = 2.05). Besides, the burden analyses showed significant differences between suicide attempters and non-suicide attempters in terms of global CNV rate and total CNV length.

So far there are two possible hypotheses for the genetic architecture of SA, i.e. the common disease common variant (CDCV) or the common disease rare variant (CDRV). Previous studies exploring CNVs for SA mainly focused on the rare and large CNVs (Perlis et al., 2012; Gross et al., 2015; Sokolowski et al., 2016b), but to date no validated CNV has been identified yet. An alternative theory for the model of complex disease is that both the CDCV and CDRV are valid, that is to say, functional variants with a range from rare to common frequencies are likely to be involved in the genetic etiology of SA (Sullivan et al., 2012). Perlis *et al.* also speculated that common CNV might contribute to SA after they could not find any rare CNVs that distinguished the cases and controls among the STAR*D subjects (Perlis et al., 2012). To our knowledge, the CNP we identified in the present study is the first reported common CNV that may be involved in the etiology of SA in major depression.

It is worthwhile to mention that the several SNPs that probe the CNP has a significant association with the expression level of *ZNF33B* in two brain regions based on the GTEx database. It is possible that the CNP might influence the risk of SA *via* affecting the expression of the gene. The function of the *ZNF33B* gene is currently not well understood. The *ZNF33B* gene, consisting of nine exons and encoding a protein of 778 amino acids, is expressed in the brain (Deloukas et al., 2004). The encoded protein belongs to the krueppel C2H2-type zinc-finger protein family. In the present study, all the top 5 associated genes in the CNV gene-based analysis (by PLINK) belonged to the zinc-finger protein family. Moreover, we searched the NCBI-GEO database and found one related study (GSE66937, public on Dec. 01, 2018) which compared the transcriptomes of suicide completers with deceased controls. After re-analyzing the gene expression profiling of the two groups, we identified three genes in this family that had significantly differential expression, including the *ZNF518A* gene (logFC = 0.31, adjusted p-value = 0.025), *ZNF254* gene (logFC = 0.25, adjusted p-value = 0.023) and *ZNF782* gene (logFC = 0.12, adjusted p-value = 0.024). In another study on bipolar disorder, epistasis between *ZNF33B* and another gene *EPHB3* on Chr. 3 showed the largest effect size (OR = 3.4) (Wang et al., 2017). Another member of this family ZNF408A was reported to be involved in neural activation and cognitive performance (Walters et al., 2010; Walter et al., 2011). ZNF804A is also one of the most promising candidate genes for schizophrenia identified in GWAS (O'Donovan et al., 2008). Taken together, the current study together with previous works suggest that zinc-finger protein family members may be involved in the genetic etiology of suicidal behavior and psychiatric disorders, and may warrant further experimental studies.

We also observed that the association between the associated CNP interacts with neuroticism in affecting the risk of SA. Furthermore, the stratified analysis revealed a greater genetic effect in subjects with a high level of the neuroticism domain (OR = 2.58) than that in the deletion groups (result not significant). These findings suggested that further studies on the interaction between genetic variants and relevant personality/psychological traits may facilitate the discovery of susceptibility variants for suicidal behavior. Indeed, it has been suggested that heritable intermediate phenotypes such as impulsivity, aggression and other relevant traits should be taken into account in the genetic study of suicidal behavior (van Heeringen, 2012; Mirkovic et al., 2016). The present study implied that personality traits such as neuroticism could be considered a promising intermediate phenotype, which may be utilized to identify a more homogeneous subgroup among heterogeneous SA subjects. With this approach, genetic variants may be discovered with a smaller but more homogeneous sample set.

So far several algorithms have been developed to detect CNVs from the SNP genotyping data (Colella et al., 2007; Wang et al., 2007), but most of these algorithms were reported to have some kinds of strengths and weaknesses including missed copy number events or no way of ranking the likelihood of such events (Winchester et al., 2009). Although the employed algorithms successfully located the position of the associated CNP from the SNP- array data, the copy number of the region for some subjects were different from the validation results using qRT-PCR. We believe our two-step approach may be useful for other CNV studies: the genome-wide CNV study *via* SNP-arrays could be used to locate the region of the associated CNVs, however one should follow with validation assays with the qRT-PCR or multiplex ligation-dependent probe amplification (MPLA) technology to determine the accurate copy number of the CNVs.

To the best of our knowledge, the current study is also the first to conduct GWAS on SA in patients with major depression in a Chinese population. Although the SNP-by-SNP association test did not reveal genome-wide significant signals, the gene-based association analysis listed *TPH2* as the top candidate gene, which is a biologically interesting candidate with literature support. TPH2, the isoform II of the rate-limiting enzyme in the biosynthesis of serotonin, encodes for the main serotonin-synthesizing enzyme in neurons. Given this functional relevance, *TPH2* gene has been considered a promising candidate in genetic association studies (Mann et al., 1997). In case-control association studies for candidate TPH2 polymorphisms, nine studies identified significant associations with SA in genotypic or haplotypic distributions, while some studies reported negative results (Lopez de Lara et al., 2007; Lopez et al., 2007; Fudalej et al., 2010; Zhang et al., 2010; Buttenschon et al., 2013). The previous two genome-wide association studies of SA in major depression did not find any signal near or located at the *TPH2* gene, although they reported other possible associations (Perlis et al., 2010; Schosser et al., 2011). Given the well-known biological function of this protein in the pathophysiology of suicidal behavior, together with the finding in this GWAS study, we believe further replication and experimental studies to elucidate the role of *TPH2* in suicide may be warranted.

We note that there are several limitations of this study and the results should be interpreted with caution. Because of the relatively small sample size, the GWAS approach and the following meta-analysis did not find any genome-wide significantly associated SNPs. The negative results from the relatively small sample size could not exclude the role of single nucleotide changes in the genetic etiology of SA in Chinese. A much larger sample or a more homogeneous group of subjects with SA are warranted in future studies. Besides, the current study did not assess the other core diathesis factors of subjects such as aggressiveness and hopelessness (Baca-Garcia et al., 2006; Jylha et al., 2016). Another important is that in general CNV calling from SNP arrays are far from perfect, as detailed in the above discussions. SNP-arrays also have limited power to detect CNV of small sizes, and the CNV region cannot be localized with a high accuracy. Recently, next-generation sequencing methods provide an alternative approach to detect CNV of smaller sizes and are better at breakpoint prediction. However, the cost is still much higher than SNP arrays especially if large sample sizes are required to detect modest-effect variants (Li and Olivier, 2013).

In summary, the present genome-wide CNV study together with validation assays identified a novel CNP within 10q11.21 that may be involved in the genetic etiology of SA in major depression, possibly by affecting the expression of *ZNF33B* in the brain. Moreover, we observed that the CNP interacted with neuroticism in affecting SA risk. These findings suggested a role of common CNV in the etiology of SA, which may open a new avenue for investigating the genetic architecture of SA.

### Ethics and Consent to Participate

Written informed consent was obtained from all the subjects after completely explaining the purpose and process of this study to them. This study was reviewed and approved by the Joint Chinese University of Hong Kong - New Territories East Cluster Clinical Research Ethics Committee (*CRE Reference No.: 2006.393*).

## Supporting information

supplementary Tables and Figures in main text

## Acknowledgements

We wish to thank all the participants and their family members, the ward staff and other associated helpers who have been very supportive of this study. Financial supports from Health and Medical Research Fund (*HMRF, Ref. No.: 12131101 and 03144526*) are acknowledged. Dr. Hon-Cheong So is supported by the Lo Kwee Seong Biomedical Research Fund and a Chinese University of Hong Kong Direct Grant.

## Contributors

Shitao RAO performed the DNA extraction and the SNP-array assay, carried out the validation assays, performed the statistical analyses and prepared the first draft of the manuscript.

Xinyu HAN and Guangming LIU gave help in the validation assays.

Marco LAM and YK WING recruited subjects as well as diagnosed and assessed psychological correlates of the subjects.

Mai SHI provided suggestions to the statistical analyses and the writing of the manuscript.

Hon-Cheong SO and Mary WAYE substantially contributed to the study design, general study coordination and the writing of the manuscript.

All authors contributed to and have approved the final manuscript.

