## supplementary Tables and Figures in main text for "Genome-wide copy number variation-, validation- and screening study implicates a novel copy number polymorphism associated with suicide attempts in major depressive disorder"

**Table S1. Demographics of the cleaned SA (n=43) and non-SA (n=43) after quality controls.**

| Phenotype | SA (n=43) | non-SA (n=43) | Statistics | P-value |
|---|---|---|---|---|
| Female (%) | 31(72.1) | 28(65.1) | $X^2$ = 0.49 | 0.486 |
| Age, mean ± SD | 42.2 ± 11.2 | 43.2 ± 13.5 | t = -0.37 | 0.710 |
| Employment (%) | 13(30.2) | 23(53.5) | $X^2$ = 4.78 | **0.029** |

SA: MDD patients with suicide attempts; non-SA: MDD patients without suicide attempts. P-value below 0.05 is in bold.

**Table S2. SA associated signals at the 10e-06 and 10e-05 levels in the pilot GWAS.**

| CHR | SNP | BP [a] | Associated allele | P-value | OR | Lower 95% CI | Upper 95% CI |
|---|---|---|---|---|---|---|---|
| 4 | rs4863321 | 1.89E+08 | A | 2.83E-06 | 16.26 | 3.703 | 71.37 |
| 2 | kgp8948717 | 47907476 | A | 3.54E-06 | 4.51 | 2.346 | 8.67 |
| 2 | rs1194480 | 65803661 | G | 8.44E-06 | 7.821 | 2.85 | 21.46 |
| 11 | rs11226619 | 1.05E+08 | A | 8.71E-06 | 0.241 | 0.127 | 0.4576 |
| 11 | rs7943892 | 1.05E+08 | C | 8.71E-06 | 0.241 | 0.127 | 0.4576 |
| 11 | rs10895780 | 1.05E+08 | A | 8.71E-06 | 0.241 | 0.127 | 0.4576 |
| 11 | rs10895781 | 1.05E+08 | C | 8.71E-06 | 0.241 | 0.127 | 0.4576 |
| 9 | kgp9618057 | 1852985 | A | 9.88E-06 | 5.148 | 2.4 | 11.05 |
| 23 | kgp22806324 | 32199287 | G | 1.53E-05 | 6.981 | 2.679 | 18.19 |
| 11 | rs11226623 | 1.05E+08 | G | 1.72E-05 | 0.2531 | 0.1335 | 0.4797 |
| 11 | rs1842893 | 1.05E+08 | G | 1.72E-05 | 0.2531 | 0.1335 | 0.4797 |
| 11 | rs1503381 | 1.05E+08 | A | 1.72E-05 | 0.2531 | 0.1335 | 0.4797 |
| 11 | kgp8455852 | 1.05E+08 | A | 1.72E-05 | 0.2531 | 0.1335 | 0.4797 |
| 11 | kgp7517932 | 1.05E+08 | A | 1.72E-05 | 0.2531 | 0.1335 | 0.4797 |
| 8 | kgp11187665 | 22074169 | G | 2.07E-05 | 10.1 | 2.903 | 35.15 |
| 4 | rs11728000 | 1.89E+08 | G | 2.08E-05 | 13.57 | 3.07 | 59.97 |
| 1 | rs6684405 | 63528577 | G | 2.67E-05 | NA | NA | NA |
| 19 | kgp1686505 | 35313251 | C | 2.67E-05 | 0 | 0 | NaN |
| 16 | rs8055870 | 72972090 | A | 2.74E-05 | 0.2012 | 0.09148 | 0.4427 |
| 22 | kgp3948966 | 37967432 | A | 2.74E-05 | 0.2012 | 0.09148 | 0.4427 |
| 16 | kgp4524138 | 72939895 | A | 2.90E-05 | 0.2596 | 0.1361 | 0.495 |
| 16 | kgp1356609 | 81673806 | G | 3.95E-05 | 0.1051 | 0.03014 | 0.3668 |
| 4 | rs2215762 | 18910602 | G | 4.37E-05 | 4.116 | 2.044 | 8.287 |
| 22 | kgp5731145 | 37965689 | G | 4.37E-05 | 0.243 | 0.1207 | 0.4892 |
| 22 | rs2235338 | 37965880 | A | 4.37E-05 | 0.243 | 0.1207 | 0.4892 |
| 22 | rs2281097 | 37966060 | G | 4.51E-05 | 0.2252 | 0.1069 | 0.4746 |
| 4 | rs7676193 | 18889088 | A | 4.57E-05 | 3.795 | 1.97 | 7.31 |
| 11 | rs2861776 | 40056052 | A | 4.77E-05 | 0.1639 | 0.06361 | 0.4222 |
| 11 | rs11035659 | 40067242 | G | 4.77E-05 | 0.1639 | 0.06361 | 0.4222 |
| 23 | rs3116883 | 1.12E+08 | A | 4.85E-05 | 0.2451 | 0.1226 | 0.4899 |
| 1 | rs6696559 | 63521891 | A | 5.04E-05 | NA | NA | NA |
| 1 | kgp4343893 | 1.9E+08 | A | 5.12E-05 | 4.732 | 2.148 | 10.42 |
| 12 | rs10876862 | 56318043 | G | 5.12E-05 | 4.732 | 2.148 | 10.42 |
| 2 | rs6742674 | 1.84E+08 | A | 5.13E-05 | 3.986 | 2.002 | 7.935 |
| 6 | rs103194 | 80569885 | A | 5.85E-05 | 0.1506 | 0.05453 | 0.416 |
| 6 | rs346305 | 80576353 | A | 5.85E-05 | 0.1506 | 0.05453 | 0.416 |
| 10 | rs1748326 | 26948937 | A | 5.93E-05 | 0.2581 | 0.131 | 0.5087 |
| 17 | kgp5953658 | 1835293 | A | 6.82E-05 | 7.484 | 2.463 | 22.74 |
| 21 | rs405227 | 24442427 | A | 6.97E-05 | 0.1835 | 0.07501 | 0.4491 |
| 8 | kgp10700041 | 1.2E+08 | G | 7.17E-05 | 0.2074 | 0.09144 | 0.4703 |

| | | | | | | | |
|---|---|---|---|---|---|---|---|
| 1 | rs487974 | 2.39E+08 | G | 7.34E-05 | 0.287 | 0.1533 | 0.5373 |
| 3 | kgp11438114 | 1.22E+08 | G | 7.60E-05 | 3.694 | 1.905 | 7.162 |
| 3 | rs1802757 | 1.22E+08 | A | 7.60E-05 | 3.694 | 1.905 | 7.162 |
| 8 | rs7004626 | 17243260 | A | 8.04E-05 | 0.2546 | 0.1264 | 0.5129 |
| 5 | kgp6766589 | 1.43E+08 | G | 8.31E-05 | 0.2363 | 0.112 | 0.4984 |
| 11 | rs4628651 | 39522637 | C | 8.31E-05 | 0.2363 | 0.112 | 0.4984 |
| 13 | kgp7960543 | 27491695 | A | 8.31E-05 | 4.232 | 2.006 | 8.927 |
| 21 | rs2831270 | 29324934 | A | 8.90E-05 | 0.1731 | 0.06702 | 0.4469 |
| 3 | kgp6076970 | 52291078 | A | 9.27E-05 | 0.2809 | 0.1468 | 0.5378 |
| 17 | rs11649913 | 14547787 | G | 9.27E-05 | 3.56 | 1.86 | 6.814 |
| 1 | rs7548511 | 8267513 | G | 9.41E-05 | 0.2629 | 0.132 | 0.5235 |
| 1 | rs11121129 | 8268095 | A | 9.41E-05 | 0.2629 | 0.132 | 0.5235 |
| 11 | rs487962 | 1.16E+08 | A | 9.41E-05 | 3.804 | 1.91 | 7.576 |
| 22 | rs16997104 | 47968920 | G | 9.41E-05 | 3.804 | 1.91 | 7.576 |
| 12 | rs1052206 | 56348028 | G | 9.43E-05 | 4.504 | 2.042 | 9.935 |
| 19 | rs918447 | 35316444 | A | 9.47E-05 | 0 | 0 | NaN |
| 19 | kgp3893752 | 35320097 | A | 9.47E-05 | 0 | 0 | NaN |
| 19 | kgp1387644 | 35320653 | A | 9.47E-05 | 0 | 0 | NaN |
| 23 | rs3126002 | 1.12E+08 | C | 9.53E-05 | 0.2605 | 0.1309 | 0.5186 |
| 1 | kgp2799241 | 37600801 | A | 9.74E-05 | 0.2016 | 0.08582 | 0.4735 |
| 12 | kgp243615 | 1.22E+08 | G | 9.74E-05 | 0.2016 | 0.08582 | 0.4735 |

[a] BP: base position (hg19); NaN: undefined; NA: not available.

**Figure S1. Chromosome position of the 3 probes in 10q11.21 from the SNP-array.** The coordinate of human genome is based on the hg38.

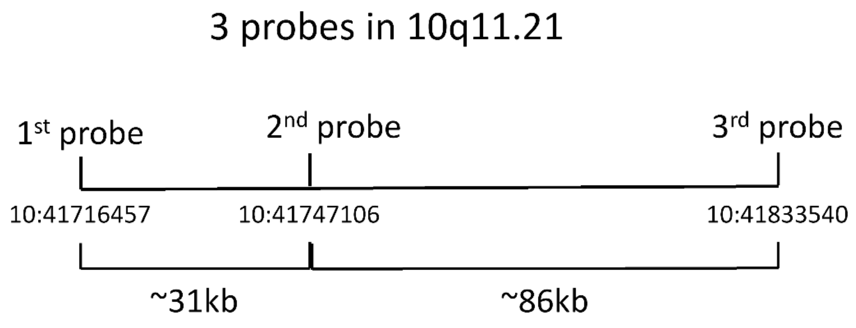

**Figure S2. SNP-by-SNP Manhattan plots for the pilot GWAS of suicide attempts within (A) the whole genome and (B) the chromosome 11 respectively.** The chromosomes are presented in order (pter to qter) and color-coded for ease of identification. The individual SNPs are represented by open circles and their corresponding p-values are graphed according to the −log10 (p-value).

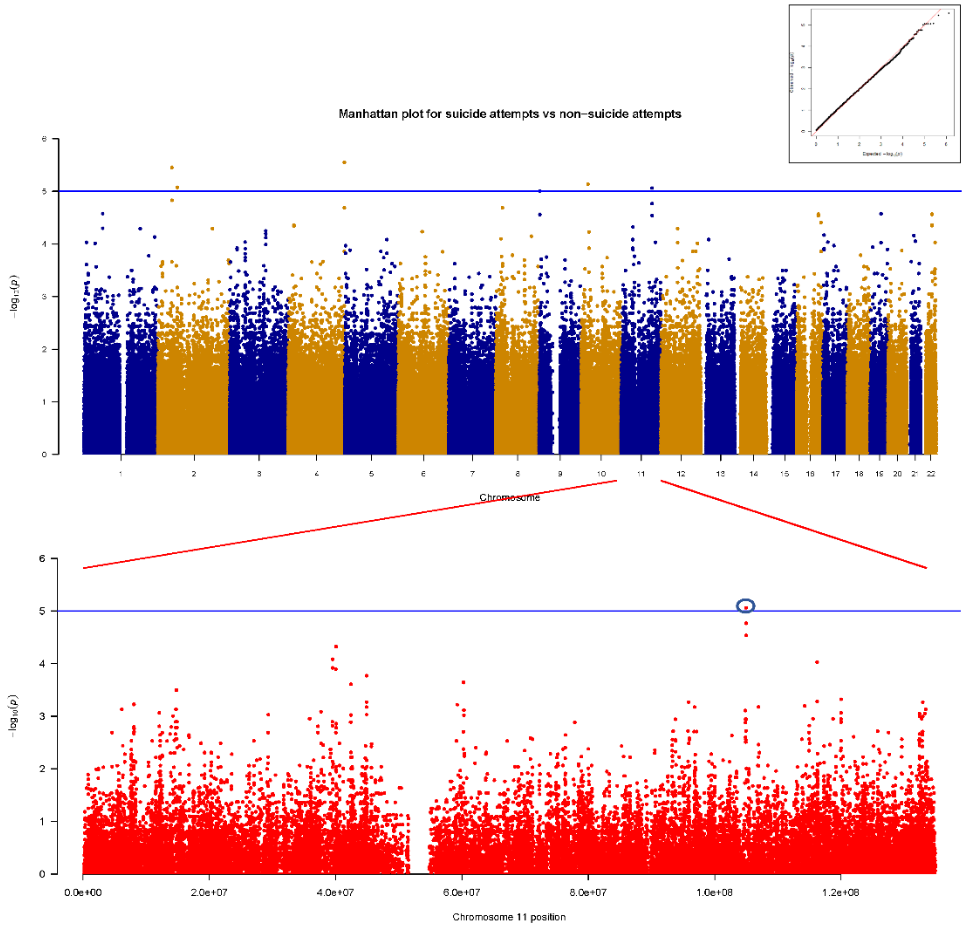

**Figure S3. Construction of LD matrix for the four top GWAS signals in Chr. 11.** Genotyping data is from the population panels of Chinese Han in Beijing and Southern Han Chinese in the 1000 genomes project database.

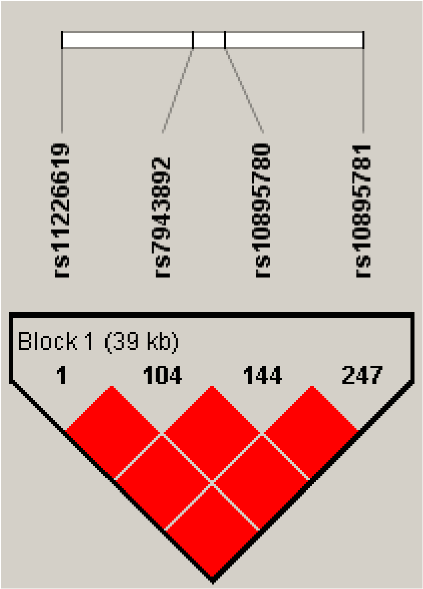

**Figure S4. Gene-based Manhattan plots for the pilot GWAS of suicide attempts.** The chromosomes are presented in order (pter to qter) and color-coded for ease of identification. The individual genes are represented by open circles and their corresponding p-values are graphed according to the −log10 (p-value).

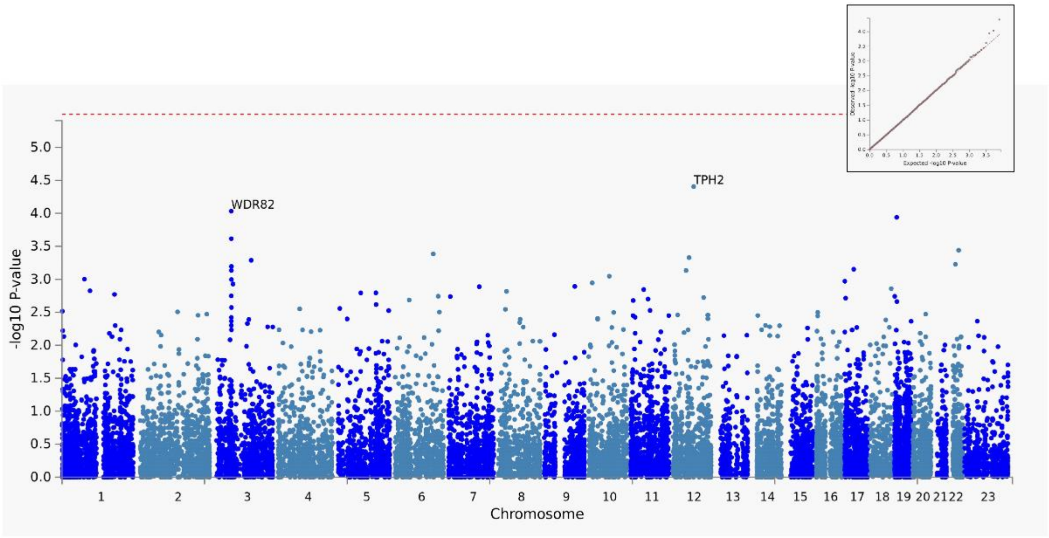

**Figure S5. CNV Value and Log R Ratio indicating presence of duplications within the region covered by the 3 probes (blue arrows) but absence of duplications in the *ZNF33B* gene (red arrows). A:** example of 3 copies duplication and normal 2 copies in the two regions respectively; **B:** example of 4 copies duplication and normal 2 copies in the two regions respectively; **C:** example of normal 2 copies in the two regions.

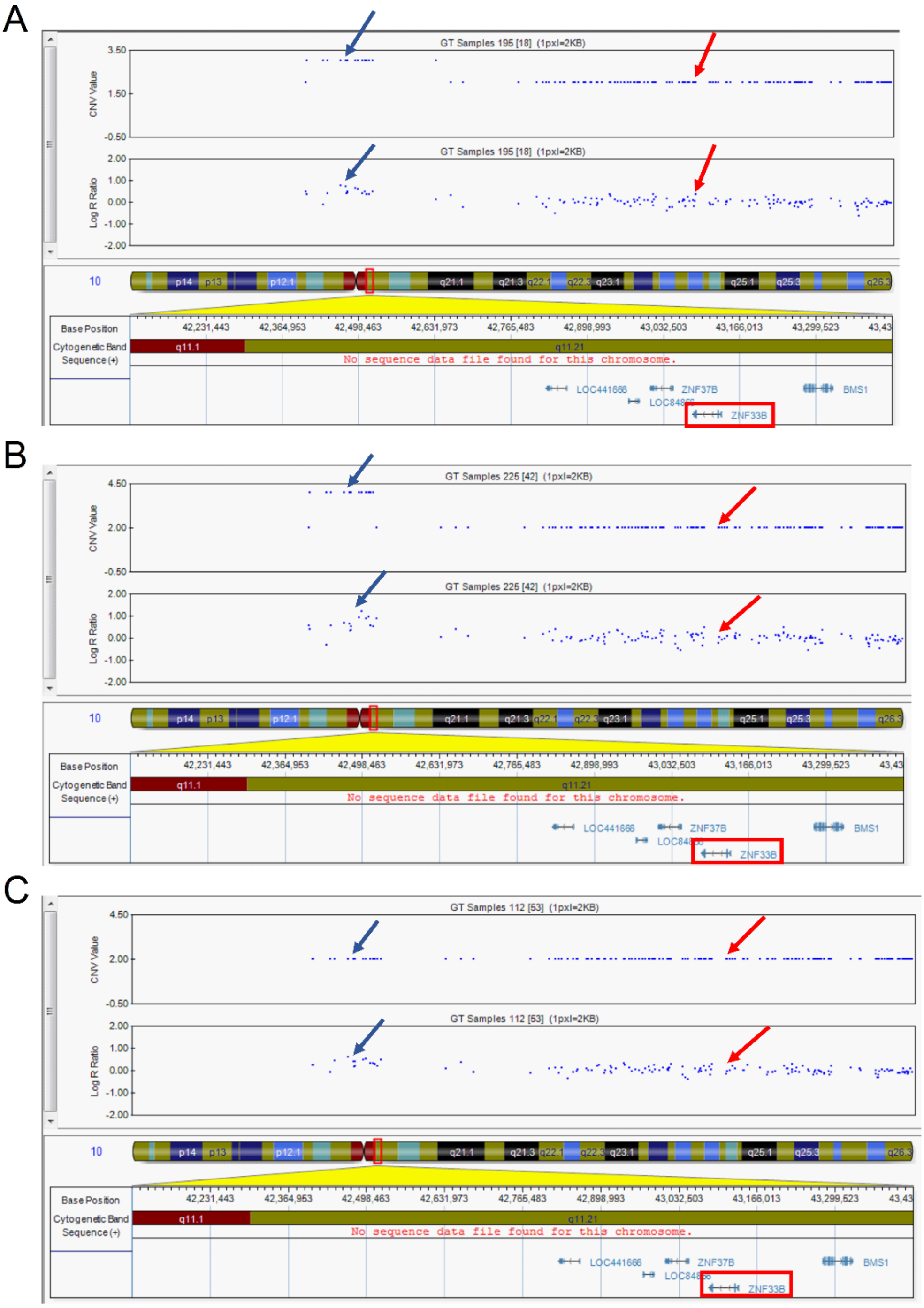

**Figure S6. Validation of the associated CNV from SNP-array data with the 1st probe (left) and the 2nd probe (right) in 9 SA (grey) and 9 non-SA (white) subjects.** Data was normalized to the RNase P endogenous control. Each subject was probed in triplicate. The calculated mean and range of copy number were presented.

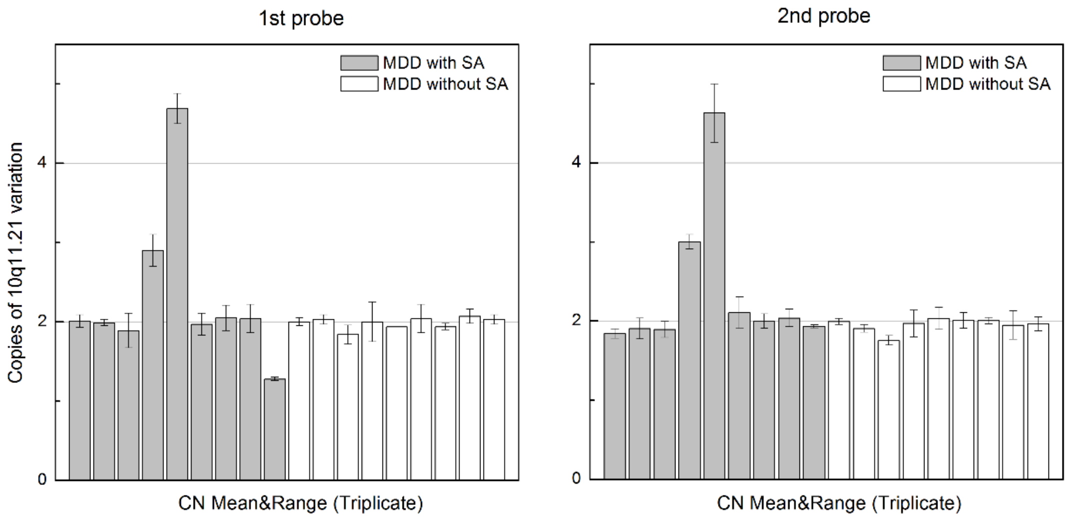

**Figure S7. Validation of the associated CNV from SNP-array data with the 3rd probe in 9 SA (grey) and 9 non-SA (white) subjects.** Data was normalized to the RNase P endogenous control. Each subject was probed in triplicate. The calculated mean and range of copy number were presented.

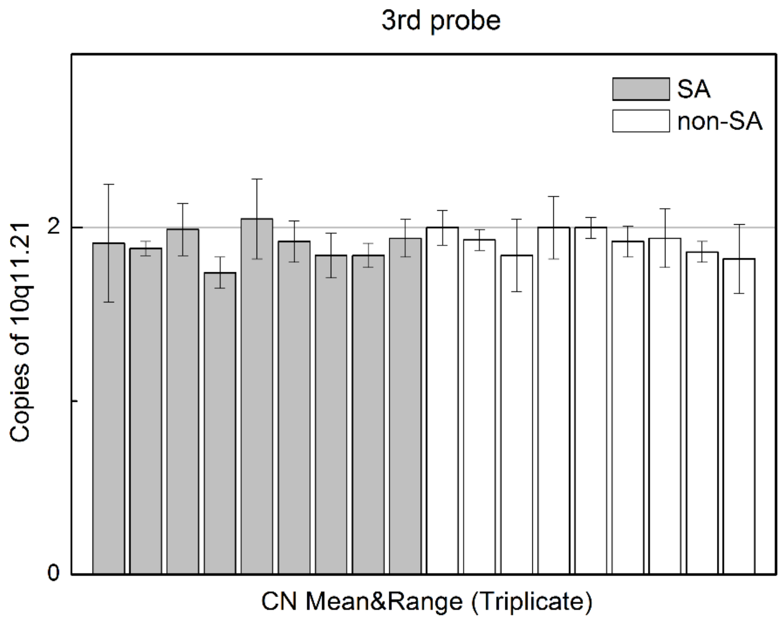

**Figure S8. Validation of the CNV within the *ZNF33B* gene from SNP-array data in 9 SA (grey) and 6 non-SA (white) subjects.** Data was normalized to the RNase P endogenous control. Each subject was probed in triplicate. The calculated mean and range of copy number were presented.

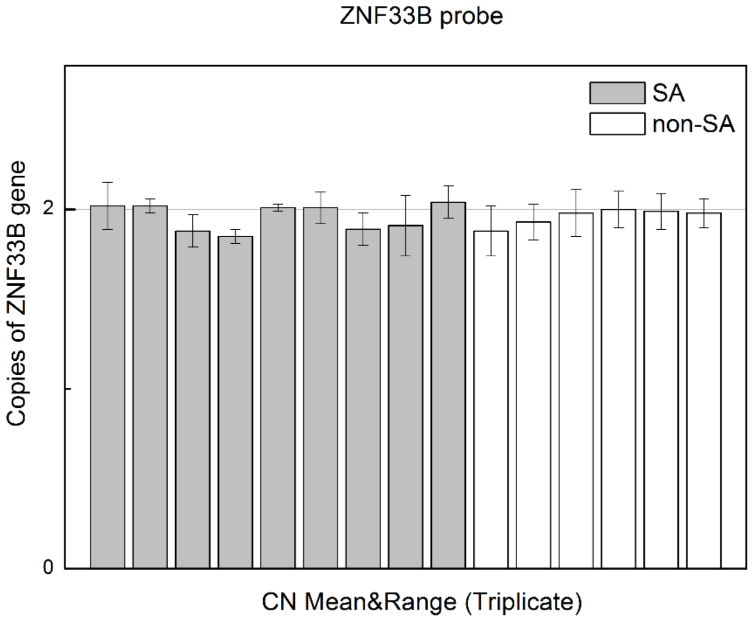
